# Classification of non-coding variants with high pathogenic impact

**DOI:** 10.1101/2021.05.03.442347

**Authors:** Lambert Moyon, Camille Berthelot, Alexandra Louis, Nga Thi Thuy Nguyen, Hugues Roest Crollius

## Abstract

Whole genome sequencing is increasingly used to diagnose medical conditions of genetic origin. While both coding and non-coding DNA variants contribute to a wide range of diseases, most patients who receive a WGS-based diagnosis today harbour a protein-coding mutation. Functional interpretation and prioritization of non-coding variants represents a persistent challenge, and disease-causing non-coding variants remain largely unidentified. Depending on the disease, WGS fails to identify a candidate variant in 20-80% of patients, severely limiting the usefulness of sequencing for personalised medicine. Here we present FINSURF, a machine-learning approach to predict the functional impact of non-coding variants in regulatory regions. FINSURF outperforms state-of-the-art methods, owing to control optimisation during training. In addition to ranking candidate variants, FINSURF also delivers diagnostic information on functional consequences of mutations. We applied FINSURF to a diverse set of 30 diseases with described causative non-coding mutations, and correctly identified the disease-causative non-coding variant within the ten top hits in 22 cases. FINSURF is implemented as an online server to as well as custom browser tracks, and provides a quick and efficient solution to prioritize candidate non-coding variants in realistic clinical settings.

## Main

Whole genome sequencing (WGS) is increasingly used to diagnose pathogenic genetic variants in patients. However, interpretation of whole genome sequencing data remains virtually restricted to the ~2% that encode proteins, because the only mutations we functionally understand well are those affecting codons and splice sites. Disease-causing non-coding variants, on the other hand, presumably operate by deregulating gene expression. Regulatory mutations causing genetic diseases are known to show a wide range of penetrance in mammals^1,22^. Still, numerous examples demonstrate that a single base pair change in a transcription factor binding site or in a gene promoter can disrupt gene expression and cause a pathology^2–6^. Identifying such regulatory variants remains eminently challenging, as it requires demonstrating that the mutation modifies gene expression timing, intensity or cell type, and leads to a disease phenotype. This may explain why large fractions of patients participating in WGS cohorts do not receive a molecular diagnosis^7,8^.

A first issue to identify candidate pathogenic variants in patients is the sheer volume of genetic variants to consider. A WGS dataset delivers 4 to 5 million variants per individual. Less than 10% are present at high frequency in the human population (more than 5% of individuals), the vast majority (>90%) are present at lower frequencies^8^, and a few thousand are unique to each genome^9^. Ultra-rare variants, among which a mutation causing a highly penetrant genetic disease might be sought in priority, therefore amount to tens of thousands of candidates. Secondly, the catalogue of regulatory elements in the human genome is still incomplete. Tremendous progress has been made in characterising the properties of non-coding regions using genome-wide assays in many cell types, including the ENCODE^10^ and the Roadmap Epigenomics project^11^. However the relationship between epigenomic signals as well as their amount of true and false positives have yet to be established^12^. Studies consistently report more than two million predicted regulatory regions in the human genome^11,13^, but most of these are not experimentally validated. Large-scale reporter assays can quantify the impact of individual variants on gene expression, but being ectopic, they do not account for the complex cognate genomic context of true regulatory regions^14,15^. Finally, linking a regulatory mutation to the gene whose expression is modified remains a critical step to demonstrate disease causality. Regulatory regions lie in vast expanses of non-coding DNA, sometimes hundreds of kilobases from their target gene. Different approaches have attempted to link regulatory elements to genes genome-wide, using correlated expression (e.g. the FANTOM project^16^), correlated chromatin states^13,17^, physical contacts with a TSS (Capture-HiC^18,19^) or evolutionary linkage (e.g. PEGASUS^20^), but the specificity of these methods is unknown.

Considering the tremendous amount, diversity and complexity of information available on chromatin states, evolutionary conservation and genome topology associated with gene regulation, a number of computational methods based on machine learning have recently been developed to identify putatively functional non-coding variants^21^ (Supplementary table 3). Their aim is to integrate the data into a single statistical framework and rank variants through a score that reflects their functional importance or regulatory potential. Current methods suffer from three main limitations. First, specificity and sensitivity are typically evaluated using data similar to the data used for model training, and it is unclear how models generalise to new variants. Second, scores are over-simplified numerical values that do not capture the rich and heterogeneous set of annotations contributing to variant selection. Third, most methods do not assign candidate regulatory variants to a predicted target gene, or resort to a naive “nearest gene” approach to do so, even though this is inaccurate in most cases^16,22,23^.

Here we describe FINSURF, a method for ranking non-coding variants in the context of human diseases. FINSURF computes a functional score predictive of the damaging nature of the mutation, but also breaks the score down into quantified contributions from all the features, making it biologically interpretable. In addition, FINSURF associates variants to one or several putatively deregulated genes when possible, which can be confronted to known genes implicated in a disease. Using a realistic setup that replicates real patient WGS in a broad range of diseases, we demonstrate that FINSURF is able to pick out the correct damaging regulatory mutations from several million variants with high accuracy and precision.

## Results

### FINSURF accurately distinguishes true regulatory variants from a variety of negative controls

We trained random forest classifiers^24^ using 880 experimentally validated, non-coding regulatory variants as a positive training set, identified in the HGMD^5^ database as Damaging Mutations (hereafter named “HGMD-DM” variants). We sampled control variants from 31 million non-coding variants with no clinical significance in the ClinVar^25^ database as negative training sets. Importantly, we observed that negative and positive variants are unequally distributed with regard to genomic features such as introns, intergenes or promoters (Fig. 1a). This probably reflects a mixture of underlying biology of regulatory variants and bias in the HGMD database. We therefore defined three negative training sets to explore the impact of genomic variant distribution on the model ability to discriminate non-coding regulatory variants. Briefly, two extreme models were built: one with ~800,000 randomly sampled negative controls (“Random”), and one with ~6,000 negative controls sampled within 1 kb of variants from the positive set (“Local”). The first model learns from large numbers of variants across the entire genome; however, the model may be overly naive, as genomic distribution biases in the training set may result in poor discrimination between closely located variants. The second model separates positive and negative variants at high resolution; but it explores a small fraction of the genome and may not generalize. We also defined an intermediate model (“Adjusted”), where we sampled ~67,000 negative controls from cytogenetic bands containing positive controls, as done by Genomiser^26^, and then sub-sampled to match the proportions of positive and negative variants within genomic feature intervals as defined by GENCODE biotypes (Methods, Figure 1a-b). During model training, we ensured that contributions from positive and negative training sets remain balanced (Methods). We assembled a compendium of 41 genomic features to deeply annotate all variants, including evolutionary sequence conservation^27–31^, biochemical composite annotations from hundreds of biological contexts^11,13,32^, sequence features, and predicted regulatory element–gene interactions^16,17,20,33^ (Supplementary Table 1; Fig. 1b).

**Figure 1.**
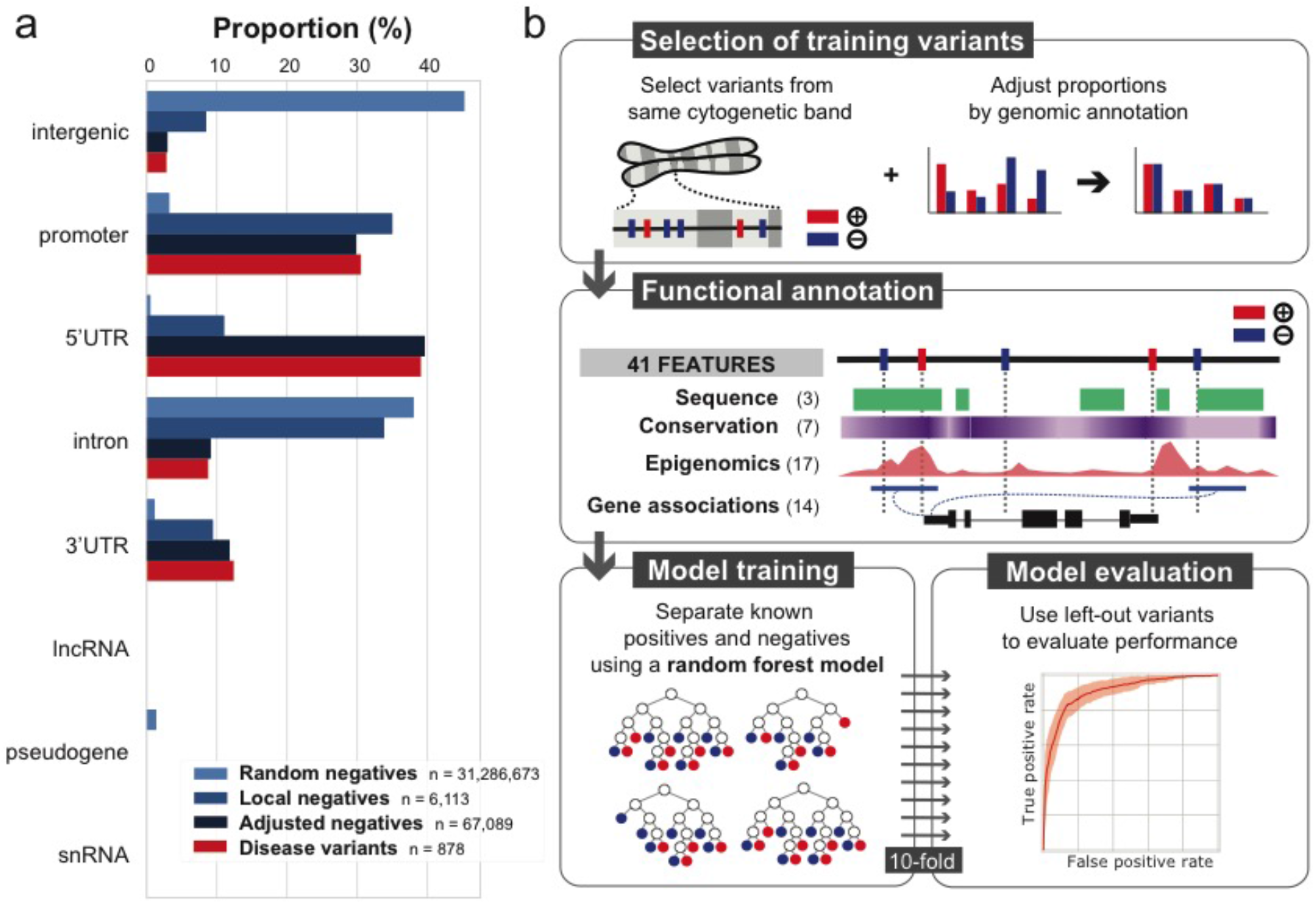
FINSURF design strategy. **a**. Percentage of genetic variants intersecting GENCODE biotypes across benign variants (shades of blue, corresponding to different sampling strategies) and damaging variants from the HGMD database (red). **b**. The final pipeline leading to the FINSURF model. Control negative variants were sampled using the Adjusted strategy. Both the negative and positive sets were annotated with 41 features, and a random forest classifier was trained to distinguish them on this basis. Ten iterations were performed, each time using 9/10 of the data, while testing performances on the remaining 1/10 which had not been used for training.

We evaluated the models abilities to distinguish regulatory from non-regulatory variants by performing 10-fold cross-validation. The positive and negative variant sets were each divided in ten subsets, and the random forest model was trained on 9 parts and its performances evaluated on the 10^th^, which contains variants not used for the training. This process was run 10 times, using each subset as the left-out evaluation subset once (Fig. 1b).

The Random model performs best, with an area under the curve (AUC) of the Receiving Operator Curve (ROC) of 0.957 and AUC of the Precision-Recall Curve (PRC) of 0.823 (Supplementary Fig. 1). Following are the Adjusted model and the Local model, in order. This performance gradient is consistent with the nature of the negative training sets. Indeed, random non-coding variants largely fall in repetitive and non-functional genomic DNA, and are easily distinguished from regulatory regions, but discrimination becomes harder as the negative set becomes more similar to the positive set. While impressive performances can be achieved by using highly contrasted negative and positive sets, such a model may perform poorly at separating the wheat from the chaff amongst closely located variants. To test this, we applied each of the three trained models to discriminate the positive variant set from the negative sets of the other two models (Supplementary Fig.1). As expected, all three models display lower performance when discriminating variants distributed differently from their own training set. Nonetheless, the Adjusted model generalises well: it performs similarly with randomly sampled negatives and its own adjusted set of negatives (ROC AUCs = 0.948 and 0.879, respectively), and just slightly underperforms compared to the Local model when using closely located positive and negative variants (ROC AUCs = 0.841 and 0.796, respectively). Because of its high performances and its ability to generalize genome-wide as well as to discriminate locally, Adjusted represents an advantageous model on which we base the rest of this study (Fig. 2a-b). We named this new model FINSURF, for Functional Interpretation of Non-coding Sequences Using Random Forests. For each variant, FINSURF provides a score, corresponding to the proportion of decision trees in the random forest classifying this variant as regulatory, as well as a description of genomic features that contributed to this classification and a list of putative target genes compiling information from its different biological features. The FINSURF score can be used to annotate, classify and rank variants relative to each other, while genomic features and target genes can be used to further refine candidate pathogenic variant searches.

**Figure 2.**
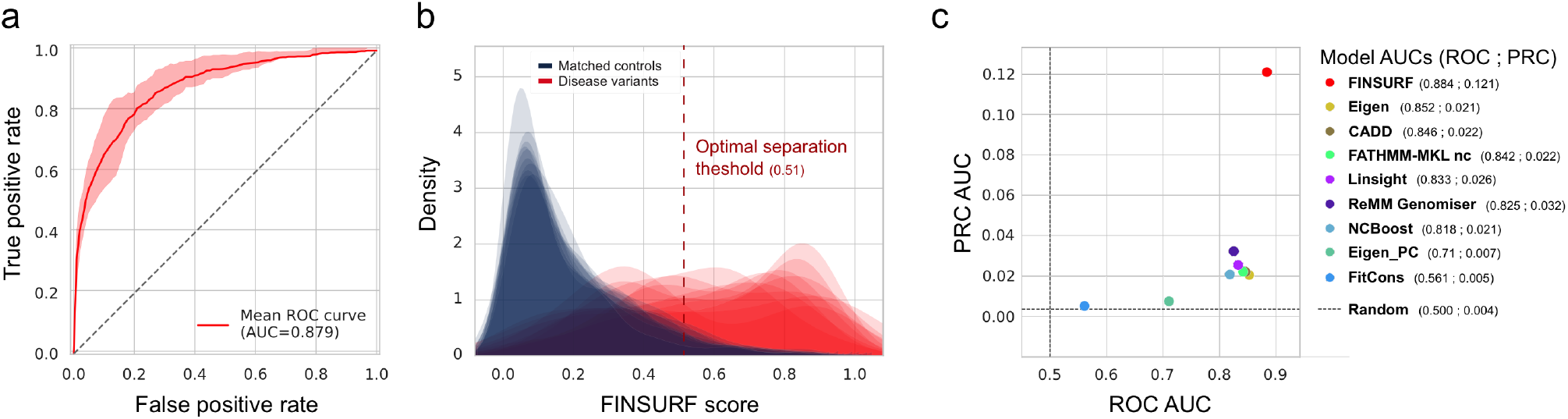
Performances of FINSURF. **a**. Receiving Operating Curve (ROC) after a 10-fold training procedure. The average curve is shown in bold red and the 95% confidence interval is indicated by a pink shading, with the mean Area Under Curve (AUC) reported in the bottom right. The dashed diagonal line indicates the distinction between positives and negatives expected by chance (AUC = 0.50). **b**. Distributions of FINSURF scores in the test set for each of the 10-fold trainings. Scores for negative variants are shown in blue, and for positive variants in red. The vertical dashed line represents the optimal score threshold (0.51) to separate positives from negatives (Methods). **c**. Scatter plot comparing classification performances (ROC AUC against PRC AUC) between FINSURF and eight other methods on a set of 62 variant independent from the training set of FINSURF. AUC values for each method and each performance metric are indicated in the legend.

### Evaluation against alternative methods and variants

We next evaluated FINSURF against eight existing methods designed to assess the functional impact of non-coding variants. For this, we performed 10-fold cross-validation again on the Adjusted set of negative variants and the set of HGMD-DM positive variants, and trained ten iterations of FINSURF. At each iteration, we also scored the evaluation subset of variants with the other methods, and we compared respective performances using ROC and PRC AUC (Supplementary Fig.2a). FINSURF outperforms all methods according to the ROC AUC values (0.819), and is second best according to the PRC AUC values (0.486) after Genomiser^26^ (PRC AUC of 0.512). Of note however, Genomiser, NCBoost^34^ and FATHMM-MKL are all trained on the HGMD-DM positive variants. Therefore these methods are placed in overly favourable conditions here, as they are evaluated on variants already used for their training, while FINSURF is evaluated on left-out variants. These results confirm that FINSURF is highly efficient to identify disease-causing, penetrant non-coding genetic variants of the type found in the HGMD-DM resource.

To verify that FINSURF’s performances extend to an independent set of non-coding regulatory mutations, we collected non-coding variants used for training Genomiser but absent from the HGMD-DM resource. This small but independent dataset comprises 92 mutations, of which 62 can be scored by all eight methods, including 41 SNV and 21 INDELS. For the negative set, 31,564 variants (of which 17,122 can be scored by all methods) were selected from the non-coding and clinically non-significative ClinVar set, following the Adjusted sampling scheme and excluding those already used to train FINSURF. Remarkably, FINSURF again outperforms all alternative methods (Fig. 2c and Supplementary Fig. 2b) despite some methods having been trained using this set of positive variants. Taken together, these results demonstrate that the combination of a carefully designed set of negative variants, a deep annotation and the optimization of a random forest classifier lead to advanced abilities to identify functional non-coding variants.

### FINSURF scores can be decomposed in biologically interpretable measurements

While accuracy is critical, most advanced statistical classification methods, including random forests, result in numerical scores, weights or probabilities that cannot be directly interpreted biologically. In addition to the score, FINSURF provides an array of descriptors for every biological feature and every variant, which serve to interpret how the model reached its conclusions. First, the Feature Importance measures how much a given biological feature contributed to discriminating positive and negative variants during training, averaged across all nodes and decision trees where the feature was sampled (Methods)^35^. Feature Importance thus provides a feature-centric view of their relative discriminatory power. Second, Feature Contributions are variant-centric measures describing how each feature individually contributed in classifying a particular variant as positive or negative^36^.

In agreement with previous models^26,34,37^, the relative Feature Importances from the FINSURF model highlight the major influence of evolutionary sequence conservation scores in classifying regulatory variants (Supplementary Fig. 3), with the Gerp^31^ score computed on a multiple alignment of 34 mammalian genomes largely dominating all other features. Additionally, the maximum predicted motif score within clusters of overlapping transcription factor binding sites (TFBS; Clustered TFBS max score) also plays a major role. Other prominent features include promoter segments and H3K27ac signals^11^, as well as enhancers from the GeneHancer^33^ collection. Together, these results confirm that FINSURF exploits diverse features consistent with different regulatory functions to identify non-coding pathogenic mutations.

Feature Contributions in turn allow users to investigate the biological properties of individual or groups of variants, and how they contributed to their classification. As an example, we relied on these variant-specific vectors to group the 880 HGMD-DM positive variants into seven clusters (Fig. 3), revealing heterogeneity in the positive training set. First, the largest cluster by far (415 variants) contains 99.8% true positives after classification by FINSURF, and is characterised by a strong evolutionary conservation signal. Second, remaining clusters with more than 50% true positives rely less consistently on sequence conservation but are all characterised by a substantial overlap with predicted TFBS clusters.

**Figure 3.**
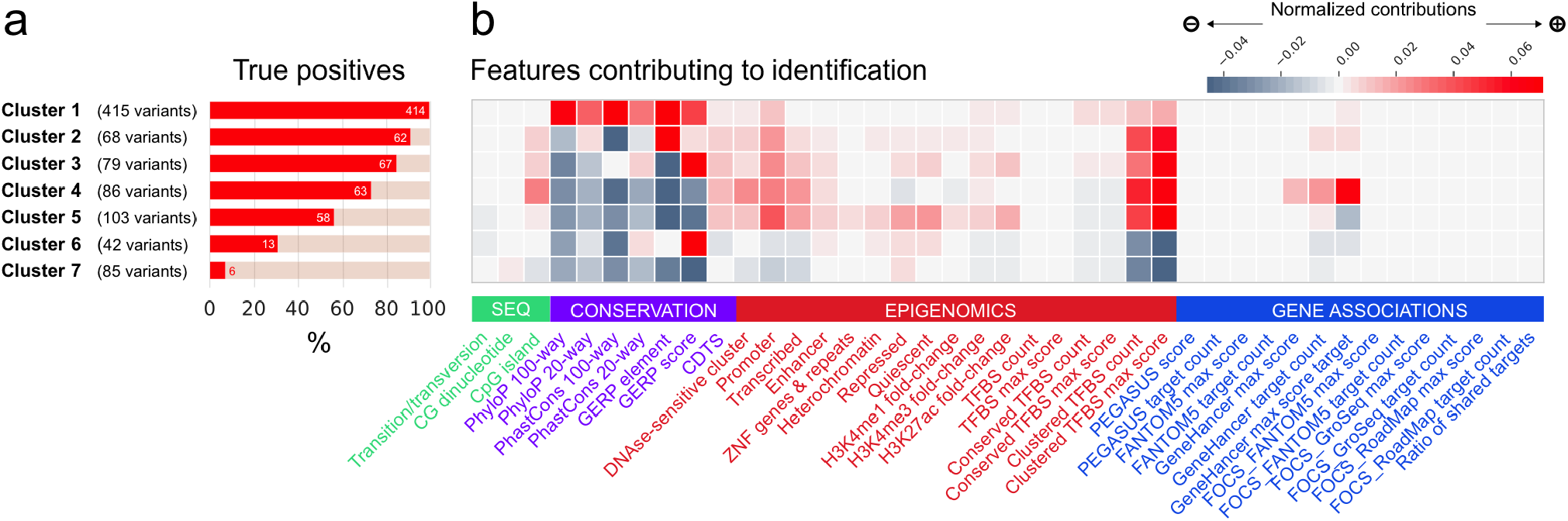
Feature contributions. **a**. The 880 positive variants were clustered using K-means into 7 clusters based on the contributions of all 41 features to their FINSURF score. Variants were classified as true positives or false negatives using the optimal score threshold (0.51). **b**. Average feature contributions in each cluster. The grey-red gradient reflects the normalized contribution of each feature and is relative across the entire grid. Features are grouped by functionally relevant categories (denoted by green, purple, red and blue colours).

Finally, clusters 6 and 7 are two small clusters where FINSURF displays low accuracy (42 and 85 variants; false negative rate > 60%). Variants in these clusters display characteristics typical of variants from the negative set and have nearly no features contributing positively to their classification. This observation could be caused by insufficient coverage of their regulatory features in the collection used by FINSURF, possibly because they are condition-dependent, or that despite manual curation of the HGMD database, these variants are in fact not regulatory.

### From whole genome sequences to pathogenic mutations

Given its high discrimination power and ability to generalize well across the genome, FINSURF is theoretically well suited to assist in diagnosing pathogenic mutations when analysing Whole Genome Sequences (WGS) from patients. To test this, we generated realistic synthetic genomes that replicate a typical clinical situation, where a patient’s genome is sequenced to diagnose the molecular cause of a known disease, but no coding mutation can be detected in any associated gene and a regulatory mutation is suspected. We focused on the set of 92 curated non-coding variants used by Genomiser for training, known to cause 56 different genetic diseases, and that were not used to train FINSURF. Despite their independence from the HGMD-DM training set, 43 variants still lied in the immediate vicinity of mutations used for training and likely share some of their biological annotations, which translated in higher but possibly overfit FINSURF scores (Supplementary Fig. 5). To alleviate this bias, we removed all variants located within 1,000 bp from any variant that FINSURF used during training, leaving 49 variants causing 30 diseases. These 49 variants display a wide distribution of FINSURF scores ranging from 0.072 to 0.965 (Fig. 4a).

**Figure 4.**
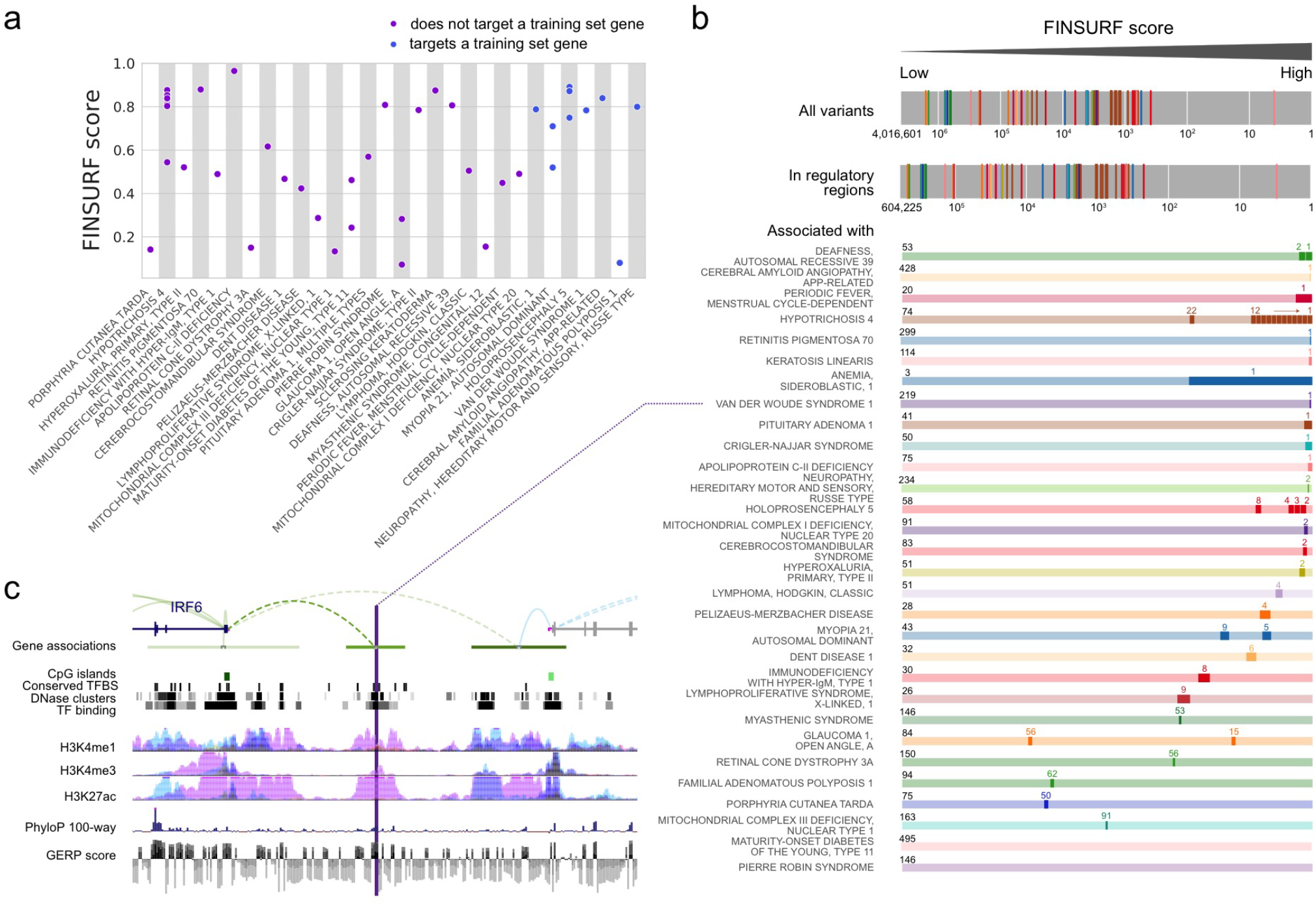
Application to medical genetics. **a**. A set of 49 regulatory variants causing human diseases (x-axis) not used for training were scored by FINSURF (y-axis). Eleven variants target a disease gene that is also targeted by a training variant (in blue), while 38 variants are totally independent (in purple). **b**. The 49 variants were seeded amongst over 4 million variants from a representative, otherwise healthy individual human genome, and their respective ranks are shown in the top bar (log scale; colors represent different diseases). When pathogenic and background variants are restricted to putatively functional non-coding sequences, the ranking does not change much (second bar). However, when filtering for regions associated with disease genes, disease-causing mutations generally show high-ranking positions (coloured bars; total number of non-coding variants associated each disease indicated on the left; pathogenic variants highlighted in dark, with their rank above). **c**. Detailed genomic context for a non-coding mutation causing van der Woude syndrome 1 (MIM 119300), which is located in an enhancer ~30 kb in 5’ to the TSS of its target gene, interferon regulatory factor 6 (IRF6). Gene associations are from the GeneHancer collection, and depict the enhancer (green horizontal bar) with the link to its predicted target gene (dashed arc). All tracks are from the UCSC genome browser.

Then, we seeded these pathogenic variants amongst 4,016,599 genetic variants identified in a reference donor from the Illumina Platinum Genome^23^ collection, and scored the resulting synthetic genome with FINSURF. Unsurprisingly, the 49 pathogenic variants rank higher than average, with the majority (82%) comprised in the top 5% (Mann-Whitney test, p value = 1.23 10^-29^, Fig. 4b). However this enrichment is of little practical use, as only one pathogenic variant scores in the top 100 variants which could realistically be investigated further by molecular biology techniques. This is an underappreciated limitation of pathogenic variant identification methods applied to real WGS data, where causative variants are vastly outnumbered by other variants, some of them with regulatory functions.

We show next how the regulatory and gene target predictions built into FINSURF dramatically increase its accuracy picking out disease-relevant regulatory candidates. We first restricted our search to variants in the 16% of the genome (471 Mb) covering predicted regulatory and evolutionary conserved regions (Supplementary Table 2). Pathogenic variants remain concentrated in the top ranking variants of this subset, with 74% in the top 5% (Mann-Whitney test, p value = 2.27 10^-23^), but still among a vast excess of false positives. Then, we relied on the OMIM database^39^ to establish lists of potentially deregulated genes for each of the 30 diseases, and selected variants within regulatory regions predicted to interact with those genes. For example, cerebral amyloid angiopathy is caused by the β-amyloid precursor protein (APP)^40^, whose regulatory regions contain 428 variants in this realistic setting. FINSURF correctly ranks the pathogenic variant in first position (Fig. 4b). Overall, across the 30 diseases included in the study, between 3 and 495 variants fall within regulatory regions targeting disease genes (average: 115). FINSURF ranks the causative pathogenic variants in first position in eleven instances. In eight other disease cases, FINSURF ranked the causative mutation in the top 5 variants, and in four additional cases, the causative mutation was in the top 10 variants. In summary, FINSURF ranked the pathogenic variant(s) within the first ten candidates for 22 out of 30 disease cases, making it the first non-coding variant predictor performing accurately in a practical, realistic setting. Of note, many of the correctly identified pathogenic variants have fairly low scores with FINSURF and other methods (Fig. 4a; Supplementary Figure 5), highlighting how target gene predictions significantly enhance the identification of relevant non-coding mutations in disease contexts.

In two disease cases, FINSURF failed to identify the pathogenic mutation because the variant lies outside of regulatory regions associated with disease gene(s). Of these, the mutation linked to Pierre Robin Sequence was initially discovered in a region located 1.4 Mb from the SOX9 gene^41^, but was later found to be present is non-affected controls and considered to be at most a modifying allele in this context^42^. FINSURF attributes a high score to this variant (0.808), but infers that this variant does not in fact regulate SOX9. Further, the two regulatory mutations associated to Maturity Onset Diabetes of the Young type 11 (MODY11) have only been shown to reduce the expression of a reporter gene *in vitro* and have no formally demonstrated role in the pathology^43^. Together, these results suggest that our methodology integrating regulatory features and target predictions shows high discrimination and accuracy to identify relevant non-coding disease variants.

## Discussion

We developed FINSURF to score variants in the human genome in the context of medical genetics, delivering an efficient strategy to identify regulatory non-coding mutations likely to cause a diagnosed disease. While a growing number of machine-learning models have been developed in recent years to identify functional non-coding variants^44,45^, few if any are applicable in practical clinical contexts, because of their design and control choices. A classical problem when designing an experiment, choices of positive and negative controls are not only paramount for model accuracy and precision, but they must also be tailored to the question that the model aims to solve. We know that a typical WGS dataset contains thousands of benign variants that occur in conserved or epigenetically active regions with characteristics of regulatory sequences^8^. Our goal is to identify a unique mutation among those that likely causes a disease. While developing FINSURF, we paid a high degree of attention to the design of a negative control dataset, and its effects on model performances and generalization. Positive controls, e.g. validated non-coding regulatory variants, remain few and far between in the literature. As a result, most models including FINSURF are trained on similarly limited datasets. However, we show here that the non-coding regulatory variants in the HGMD database are quite heterogeneous in terms of genomic location and biological features (Fig. 3), suggesting that FINSURF can capture a varied range of functional variants. Possibly less appreciated, negative controls are also crucial to developing a successful model. Randomly chosen benign human variants, or solely based on population frequency, result in models that successfully discriminate negatives from positives but lack precision within broad regulatory regions. On the other hand, benign variants closely matched to positive controls achieve excellent local discrimination but result in overly specific models. We note that previous models have been trained on sometimes elaborately sampled negative variants^26,46^, but the theoretical justifications and the consequences of those choices on model performance have rarely been explored. These considerations have been eclipsed by over-reliance on ROC curves to assess performance, but ought to be properly addressed. FINSURF was explicitly tested both at the general and the local level using different sets of controls to ensure high performances genome-wide yet serve the purpose of identifying pathogenic mutations from a background of variants in similar genomic contexts.

Nonetheless, the concept of regulatory sequence covers multiple situations. From developmental enhancers with strong effects on gene expression and organism fitness to redundant shadow enhancers, ultimately the regulatory potential of a genomic sequence is likely to be a continuous property rather than a binary characteristic. This is consistent with the distribution of FINSURF scores observed during performance tests (Fig. 2) where 6.4% of benign variants obtain scores above the optimal separation threshold of 0.51, while 41.4% of pathogenic mutations are below this threshold and therefore not distinguishable from benign variants. Solely relying on a score and threshold to identify relevant regulatory mutations from whole-genome sequences is probably bound to fail in the vast majority of medical genomics applications. To alleviate this issue, we provide an efficient strategy to first prioritize context-relevant candidate variants, and then characterise the individual functional profiles of each candidate identified by the model for further interpretation. Combining information from FINSURF scores and disease aetiology, we are able to correctly identify the causative regulatory mutation as a top candidate in an array of 23 pathologies representative of the state-of-the-art on non-coding pathogenic mutations. While current sets of such mutations arguably suffer from ascertainment bias, FINSURF may significantly broaden the pool of relevant candidates for future clinical analyses compared to prevailing approaches where exon-proximal variants are prioritized.

### Availability

FINSURF is available as an online application at https://www.finsurf.bio.ens.psl.eu/ where users can load their variants in VCF format as well as optional lists of relevant target genes. The webserver returns an interactive list of ranked candidate non-coding variants, and allows users to investigate their functional characteristics through Feature Importance graphs and links to custom tracks in the UCSC web browser. A comprehensive list of 471,099,210 non-coding positions in the human genome with relevant functional features, their FINSURF scores, and their Feature Importance values in the model, is also available in BED format for more computationally intensive projects at https://www.opendata.bio.ens.psl.eu/finsurf/. The code necessary to install FINSURF locally, to produce figures 2, 3 and 4 and to train the model is available at https://github.com/DyogenIBENS/FINSURF/.

## Methods

### Training datasets

As positive controls, we selected all variants labelled as Damaging Mutations from the Human Gene Mutation Database^5^ (version Pro 2017.2). To focus on non-coding variants, we excluded variants that were annotated as protein-impacting and/or located within CDS according to the HGMD gene annotations. We additionally used the GENCODE^47^ genome annotation (version 29 lift-over hg19) to further exclude variants located within CDS, start codon, stop codon, or splice sites. This lead to the identification of 880 non-coding damaging mutations. This entire set was included for the “Random” model, but reduced to 878 for the “Adjusted” model and to 877 in the “Local” model, as some variants did not have any appropriate negative controls (see hereafter for the description of these models and sampling schemes). Negative controls were sampled from a set of 38,056,330 variants for which no medical impact was found (accessed on 2017/09/05 from ftp://ftp.ncbi.nlm.nih.gov/pub/clinvar/vcf_GRCh37/archive_1.0/2017/).

A first set of 31,516,128 negative controls (here-after named “Random”) was sampled randomly after removal of coding variants and of indels, as no indels were present in the positive set. This corresponds to a naive approach under which no particular bias is expected between the positive and negative controls aside from the differences in distributions of annotations. A second set of 6,113 negative controls was selected within 1,000 bp of positive control variants (here-after named “Local”). This control set was built to test separation of positive and negative controls at high genetic resolution. The third set (called “Adjusted”, which was used for the analysis) corresponds to a intermediate situation, where negative controls are sampled from cytogenetic bands containing at least one positive control to avoid artificially biasing negative controls towards non-functional genomic regions^26^. Additionally, proportions of negative controls in the different GENCODE biotypes were matched to those of positive controls. This correction aims at forcing the classification model to focus on the functional differences between positive and negative controls, rather than capturing differences that arise from a location bias (as positive controls are biased towards gene-proximal non-coding regions). In total, 67,089 negative controls were retained in this set.

### Annotation of non-coding variants

See Supplementary Table 1 for a complete list of annotations. To build the classification model, we identified four groups of annotations to characterize non-coding variants. The first group corresponds to annotations related to sequence evolutionary conservation. PhyloP and PhastCons scores for the human genome (hg19 version) based on the 100 vertebrate multiple genome alignment were downloaded from the UCSC web browser. The same scores were obtained for the 20 primate genomes alignments for the hg38 version, and converted to hg19 coordinates using liftOver^48^. In addition, GERP scores and GERP elements were downloaded from the Sidow lab webpage (hg19 version). Finally, the Context-dependent tolerance score (CDTS) evaluates constraint at the human population level. Scores were obtained from the original publication^30^ for hg19. The second group corresponds to annotations describing biochemical properties of the genome. The Roadmap Epigenomics project provides with 18 chromatin states inferred from combinations of histone modifications, across 98 cell types. This large amount of features would correspond to a very sparse set of annotations, a variant being associated to a single of the 18 chromatin states for a given cell type. As sparse annotations are poorly exploited by random forests, we aggregated these chromatin states per genomic position and counted the number of cell types corresponding to each chromatin state. We also selected three key histone marks of regulatory regions, H3K4me1, H3K4me3, and H3K27ac, and calculated the median Fold Change value at each genomic position across the same 98 cell types. Additionally, we downloaded two datasets related to transcription factor binding sites from the UCSC Genome Browser: TFBS identified as conserved between mouse, rat, and human; and clusters of TFBS identified in ChIP-seq peaks from 161 experiments across 91 cell types from the ENCODE project. A complementary dataset of TFBS from JASPAR, identified within ChIP-seq peaks, was obtained from Ensembl^49^. Finally, a dataset of DNaseI hypersensitive regions was obtained from the UCSC, corresponding to clusters identified in 125 cell types from ENCODE. A third group of features describe sequence properties related to variant locations. Three features were retained: CpG dinucleotides and CpG islands, downloaded from the UCSC, and variant type (transition, transversion, or INDEL). The last group gathers annotations from recently published datasets predicting regulatory regions in the genome together with their gene targets. We extracted predicted regulatory regions from the following datasets: GeneHancer^33^ (version 4.4, accessed 2018-12-17), PEGASUS^20^ (2018-12-17), FANTOM5 co-expressed regions^16^ (2018-11-19), FOCS FANTOM5, FOCS ROADMAP DHS, and FOCS Gro-seq co-expression^17^ (2018-12-11). We also included promoters (2kb upstream of transcription start sites) and UTR regions from all coding and non-coding genes from the GENCODE^47^ dataset as potential non-coding regions of interest (v29lifthg19). These datasets were integrated by merging overlapping elements into regions with one or multiple evidences of predicted associations. This integration was not performed for GeneHancer, which was handled separately, as it does not provide a single score for each regulatory region/predicted target pair. A list of predicted targets was derived for each regulatory region by merging all predicted target genes across methodologies. In addition to these four groups of annotations, which are used for classification by the FINSURF model, other annotations were included for filtering and characterizing variants. Notably, we used the GENCODE (v29liftHg19) annotations for the locations of biotypes such as CDS, introns, etc. A total of 471,099,210 genomic positions were thus annotated with the set of descriptors, and evaluated for functional potential with FINSURF. For each position, 2 predictions are reported: one for transitions and one for transversions. Tabix-indexed tabular files for all chromosomes were generated, allowing the fast interrogation of these files for millions of variants of interest. Functional profiles are reported for all variants lying within annotated regulatory regions. For indels, all positions affected by the variant (as well as the one preceding and following positions) are evaluated, but only the highest score is reported.

### Model training and performance evaluation

We trained three Random Forest models (Random, Adjusted and Local, using different negative control sets) using the Scikit-Learn Python library v.0.20.2, with the following parameters: 1,000 trees, maximum depth = 15 nodes, minimum number of samples per leaf = 1. These parameters were optimized using a nested-cross validation evaluation of 400 models, exploring different range of values. Class size imbalance issues in the training set were solved by sampling with replacement a set of n random variants for each class, where n is the size of the smallest class (in this case, HGMD-DM non-coding variants; n = 880). Each tree in the Random Forest was then built from a different, balanced set of positive and negative controls. Model performance was evaluated using 10-fold cross-validation. We note that genetic variants in the training set are not necessarily independent, as they can be located at close genomic distance and thus share some of their features. This can lead to performance over-estimation when closely located variants of the same class are split into training and validation sets, which can artificially favour correct classification of the validation variants. To mitigate this problem, for cross-validation variants were separated by location, based on cytogenetic bands. Model discrimination between variant classes was evaluated based on the Receiver-Operating Curve (ROC; true positive rate as a function of false positive rate) and the Precision-Recall Curve (PRC; proportion of true positives among all positives, as a function of the true positive rate). This second curve is of particular interest in the context of imbalanced learning, as it better captures how the proportion of true positives against false positives changes with increasingly lenient thresholds on the prediction score. For the Adjusted model, we maximized the F1-score, defined as the harmonic mean of the precision and recall, and obtained an optimal prediction score threshold of 0.51. This threshold was used to calculate the confusion matrix. Eight other methods were also evaluated using 10-fold cross-validations on the control dataset. Variants missing a score for any of the methods were excluded from the evaluation (average drop-out rate = 53%). Indels were scored as with FINSURF: all bases covered by the indel were scored, but only the highest scoring position was retained.

Finally, we applied each of the three FINSURF models (Random, Adjusted, Local) on the training datasets of the two other models, in order to evaluate the performance of each model across different genomic contexts. We re-used the 10-fold cross-validation scheme used for model training, but evaluated the 10 partial models using negative test sets sampled with either of the other two models. In order to make the performance curves comparable, an additional random sub-sampling was performed on the negative controls, so that the proportion of positives in the test-subset of each k-fold was as close as the one from the “Local” selection scheme (12.5%).

### Independent evaluation

We downloaded 448 disease-causing non-coding variants used to train the Genomizer model^26^. Of these, we excluded 11 variants found within coding regions according to GENCODE v29. We then intersected these variants with our training set of regulatory HGMD-DM variants and excluded overlaps. This resulted in a set of 92 non-coding regulatory variants independent from our training positive controls, which were used to compute ROC and PRC curves. Of these, 30 were dropped during the comparison against other methods, as some models did not provide scores for these positions, leaving 62 variants for the comparison. Negative controls (N=17,122) were sampled from the ClinVar dataset, using the Adjusted sampling protocol. For analysis of a realistic genome-wide VCF (Figure 4), we further excluded disease-causing non-coding variants located within 1,000 bp of an HGMD-DM variant from the FINSURF training dataset, and specifically focused on 49 variants that represent a set of fully independent regulatory variants with respect to our model. All non-coding variants (according to GENCODE v29) from the Illumina Platinum genome NA12877^38^ were annotated and scored with FINSURF as a realistic genetic background.

### Feature contributions and clustering

Feature contributions were calculated using a more computationally efficient, in-house re-implementation of the treeinterpreter package (https://github.com/andosa/treeinterpreter; https://arxiv.org/abs/1906.10845), which calculates the average decrease in Gini impurity index across all trees for each feature/variant combination. To avoid overfitting, feature contributions for a particular variant were calculated only from trees where this variant was not used for training. K-means clustering was performed on the feature contributions vectors for the positive control variants to explore structure in the training set. Different K values from 2 to 19 were explored, and the optimal K was selected using maximization of the silhouette score and inertia minimization with Scikit-Learn v.0.20.2. For each cluster, mean feature contributions were calculated to obtain the average functional profile. To translate these feature contributions into feature values, we calculated the feature effect size for variants within a cluster against variants from other clusters combined with negative controls. This comparison highlights distinctions between positive and negative controls that are specific to this cluster. Effect size was calculated depending on feature distribution: Cohen’s h for binary features, and Cohen’s d for discrete and continuous features:

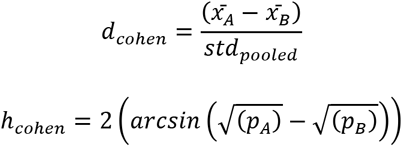

### Disease association analysis

We collected sets of genes from OMIM^39^, associated with 30 diseases caused by 49 fully-independent non-coding variants from the Genomizer^26^ training dataset (see ‘Independent evaluation’ above). Each disease was associated with a single gene, except for “Cerebral Amyloid Angiopathy, APP-related” (OMIM:605714) for which 3 genes were retrieved from the OMIM webpage. Variants within regulatory regions with predicted targets that contained the identified disease-gene were selected and ranked based on their FINSURF score.

## Acknowledgements

We wish to thank Pierre Vincens for assistance with computational infrastructure. This work was supported by the French Government and implemented by ANR (ANR-10-LABX-54 MEMOLIFE and ANR-10-IDEX-0001- 02 PSL* Research University), by the French Minister for research and Education and by the Fondation pour la Recherche Médicale, grant number FDT201805005782 to LM.

## Author contributions

LM performed analyses; LM and HRC designed the FINSURF model; AL and TTN implemented the online server; CB, LM and HRC analysed the data and wrote the manuscript.

