## Supplementary Material for "Classification of non-coding variants with high pathogenic impact"

| Feature | Description | Group | Distribution | Reference |
|---|---|---|---|---|
| Transition/Transversion | Nature of the variants (0 for transitions, 1 for transversions ; indels are set to 2). | Sequence properties | Discrete | This study |
| CG dinucleotide | CG dinucleotide locations. | Sequence properties | Binary | This study |
| CpG islands | CpG islands from the UCSC. | Sequence properties | Binary | [1] |
| PhyloP 100-way | PhyloP score computed from 100 vertebrate genomes' alignments | Conservation | Continuous | [1,2] |
| PhyloP 20-way | PhyloP score computed from 20 primate genomes' alignments | Conservation | Continuous | [1,2] |
| PhastCons 100-way | PhastCons score computed from 100 vertebrate genomes' alignments | Conservation | Continuous | [1,3] |
| PhastCons 20-way | PhastCons score computed from 20 primate genomes' alignments | Conservation | Continuous | [1,3] |
| CDTS | Context-dependent tolerance score representing the tolerance to genetic variation, evaluated at the population level. | Conservation | Continuous | [4] |
| GERP score | GERP score | Conservation | Continuous | [5] |
| GERP element | Regions identified as conserved element. | Conservation | Binary | [5] |
| DNase-sensitive cluster | Score associated to DNaseI Hypersensitivity Clusters in 125 cell types from ENCODE. | Biochemical evidence | Continuous | [6] |
| Promoter | Number of the considered state at given location across 98 cell types from Roadmap Epigenomics. | Biochemical evidence | Discrete | [7] |
| Transcribed | | Biochemical evidence | Discrete | [7] |
| Enhancer | | Biochemical evidence | Discrete | [7] |
| ZNF Genes & repeats | | Biochemical evidence | Discrete | [7] |
| Heterochromatin | | Biochemical evidence | Discrete | [7] |
| Repressed | | Biochemical evidence | Discrete | [7] |
| Quiescent | | Biochemical evidence | Continuous | [7] |
| H3K4me1 fold-change | Median of Fold Change value per position for the considered histone mark across 98 cell types from Roadmap Epigenomics. | Biochemical evidence | Continuous | [7] |
| H3K4me3 fold-change | | Biochemical evidence | Continuous | [7] |
| H3K27ac fold-change | | Biochemical evidence | Continuous | [7] |
| TFBS count | From predicted TFBS overlapped by a variant, two information are extracted : the number of predicted TFBS, and the maximum score of site prediction. | Biochemical evidence | Discrete | [8] |
| TFBS max score | | Biochemical evidence | Continuous | [8] |
| Conserved TFBS count | | Biochemical evidence | Discrete | UCSC track |
| Conserved TFBS max score | | Biochemical evidence | Continuous | UCSC track |
| Clustered TFBS count | | Biochemical evidence | Discrete | UCSC track |
| Clustered TFBS max score | | Biochemical evidence | Continuous | UCSC track |
| PEGASUS score | From predicted regulatory regions overlapped by a variant, two information are extracted : the score associated to the regulatory region, and the number of targets. When multiple regions from a dataset overlap, the maximum score is retrieved.<br><br>GeneHancer provides with an additional score for the predictions of interactions, which is extracted as "max score targets". | Gene associations | Continuous | [9] |
| PEGASUS target count | | Gene associations | Discrete | [9] |
| FANTOM5 max score | | Gene associations | Continuous | [10] |
| FANTOM5 target count | | Gene associations | Discrete | [10] |
| GeneHancer max score | | Gene associations | Continuous | [11] |
| GeneHancer target count | | Gene associations | Discrete | [11] |
| GeneHancer max score targets | | Gene associations | Continuous | [11] |
| FOCS_FANTOM5 max score | | Gene associations | Continuous | [12] |
| FOCS_FANTOM5 target count | | Gene associations | Discrete | [12] |
| FOCS_GroSeq max score | | Gene associations | Continuous | [12] |
| FOCS_GroSeq target count | | Gene associations | Discrete | [12] |
| FOCS_Roadmap max score | | Gene associations | Continuous | [12] |
| FOCS_Roadmap target count | | Gene associations | Discrete | [12] |
| Ratio shared targets | At positions overlapped by regulatory regions of from multiple source, we divide the number of genes predicted as a target by more than 1 source, by the total number of target genes across sources. | Gene associations | Continuous | This study |

**Supplementary Table 1: Sources of annotations**

| Source | Description | Number of bases | Reference |
|---|---|---|---|
| GENCODE v29liftHg19 | Extracted 5'UTR and 3'UTR regions, as well as proximal regulatory regions set as the 2,000bp before the first position of annotated genes (both coding and non-coding). | 128,404,642 | [13] |
| PEGASUS | Predicted regulatory regions identified from conserved non-coding elements. | 68,593,981 | [9] |
| FANTOM5 | Robust enhancer-promoter associations based on correlation of expression levels measured by CAGE. | 8,444,398 | [10] |
| GeneHancer | Unified predicted regulatory regions from a set of 4 sources : Ensembl regulatory built, FANTOM5 permissive enhancers, VISTA enhancers, and ENCODE proximal and distal enhancers. | 312,286,188 | [11] |
| FOCS FANTOM5 | Predictions through the FOCS pipeline of regulatory associations from FANTOM5 expressed regions. | 5,131,199 | [12] |
| FOCS GroSeq | Predictions through the FOCS pipeline of regulatory associations from global run-on sequencing datasets. | 21,636,242 | [12] |
| FOCS Roadmap | Predictions through the FOCS pipeline of regulatory associations from DHS datasets from Roadmap Epigenomics. | 13,401,492 | [12] |

**Supplementary Table 2: Sources of genomic regions integrated into the Regulome scored by FINSURF**

| Name | Model | Training scheme | Evaluation | Ref |
|---|---|---|---|---|
| Genomiser | Supervised classification with an ensemble of random forests. | Manual curation of a set of functional variants (note : a large part overlaps with HGMD), against nearly fixed human-specific variants identified within the primate lineage, taken as negative controls. | | 14 |
| NCBoost | Supervised classification with gradient-boosting of decision trees (XGBoost). | | | 15 |
| FATHMM-MKL | Supervised classification with a Support Vector Machine using a Multi-Kernel learning approach. | Positive controls are taken from HGMD. Negative controls are taken from the 1,000 genome project : variants with mAF>1 %, at a maximum distance of 1kb of a positive control. | | 16 |
| Eigen | Unsupervised learning | | | 17 |
| Linsight | | | | 18 |
| FitCons | Evaluation of sequence-constraint at base-level within different genomic regions identified from biochemical features. | | | 19 |
| CADD | Supervised classification with a Support Vector Machine using a linear kernel. | | | 20 |

**Supplementary Table 3: Variant scoring methods compared to FINSURF**

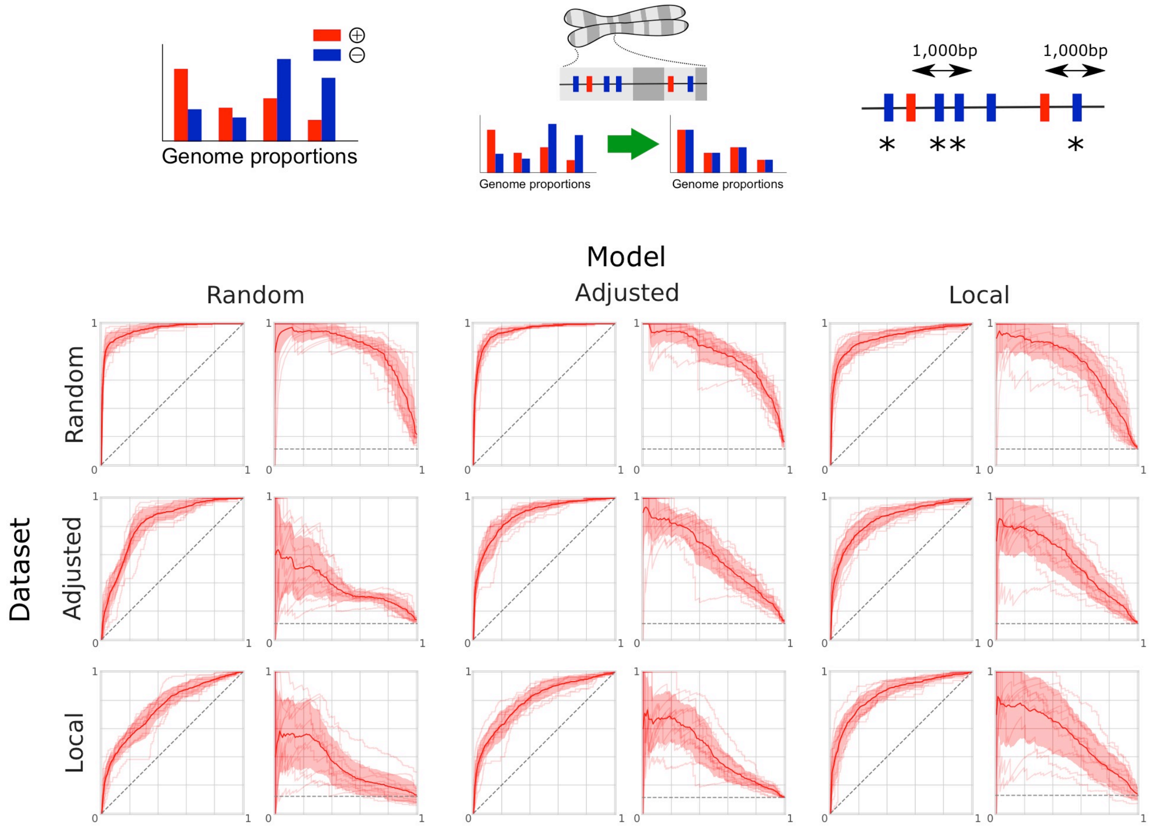


| ROC AUC (Dataset \ Model) | Random | Adjusted | Local |
|---|---|---|---|
| Random | 0.957 | 0.948 | 0.902 |
| Adjusted | 0.821 | 0.879 | 0.853 |
| Local | 0.748 | 0.796 | 0.841 |

| PRC AUC (Dataset \ Model) | Random | Adjusted | Local |
|---|---|---|---|
| Random | 0.823 | 0.773 | 0.708 |
| Adjusted | 0.375 | 0.587 | 0.552 |
| Local | 0.342 | 0.427 | 0.538 |

**Supplementary Figure 1.** Design and cross-performance evaluation of the random forests models trained with the three approaches for sampling negative control variants. As described in the methods, the Random sampling does not correct for the differences in proportions between positive and negative controls in the different genomic annotations, while the Adjusted model corrects for this. The cross-performance of the models is evaluated by comparison of the ROC and precision-recall curves, recalculated from the 10 fold cross validation step of each model (which correspond to the pairs of curves on the diagonal). A given column corresponds to the application of a model on the 10-fold subsets of variants obtained under the different samples methods (with filtering of overlapping variants between training and validation subsets, as described in the methods). Note that to allow the comparison of performances, a second undersampling of negative control variants was performed in the validation subset for the Random and Adjusted models, in order to match the observed proportion of 12.5% of positive controls in the Local sampling dataset. The area under the curves of the average curves is reported in the two tables for each combination of model and validation dataset.

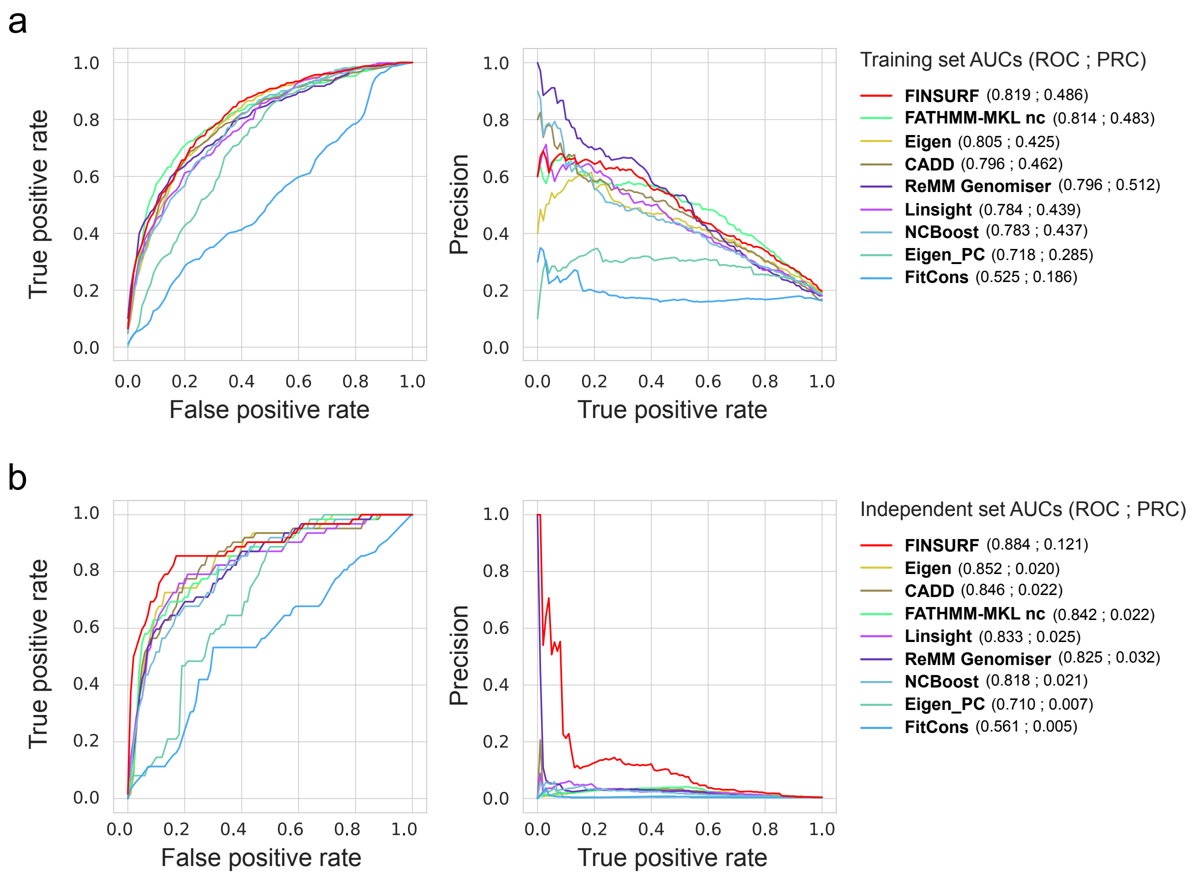


**Supplementary Figure 2.** Comparison between FINSURF (red lines) and eight other methods also designed to functionally interpret non-coding regulatory variants. **a**. Comparison of performances on the training dataset of FINSURF "Adjusted". The subsets of variants generated during the 10-fold cross-validation were re-used, by scoring the positive and negative control variants with the other methods. At no time was FINSURF tested on variants used during training. Variants for which a score could not be computed by any one method (e.g. lack of required annotations) were discarded for all methods. The average performance curve of the 10-folds are reported. We note the high precision of the ReMM Genomiser model, which can be explained by the overlap between the training variants of this model and the HGMD variants used by FINSURF. **b**. Comparison of performances on the subset of 92 Genomiser variants, of which 30 are discarded due to missing scores by at least one method. Negative control variants were re-sampled following the Adjusted sampling procedure, and those found in the training dataset of the FINSURF model were removed. Note that the ROC curve is the one presented in the main figure 2c.

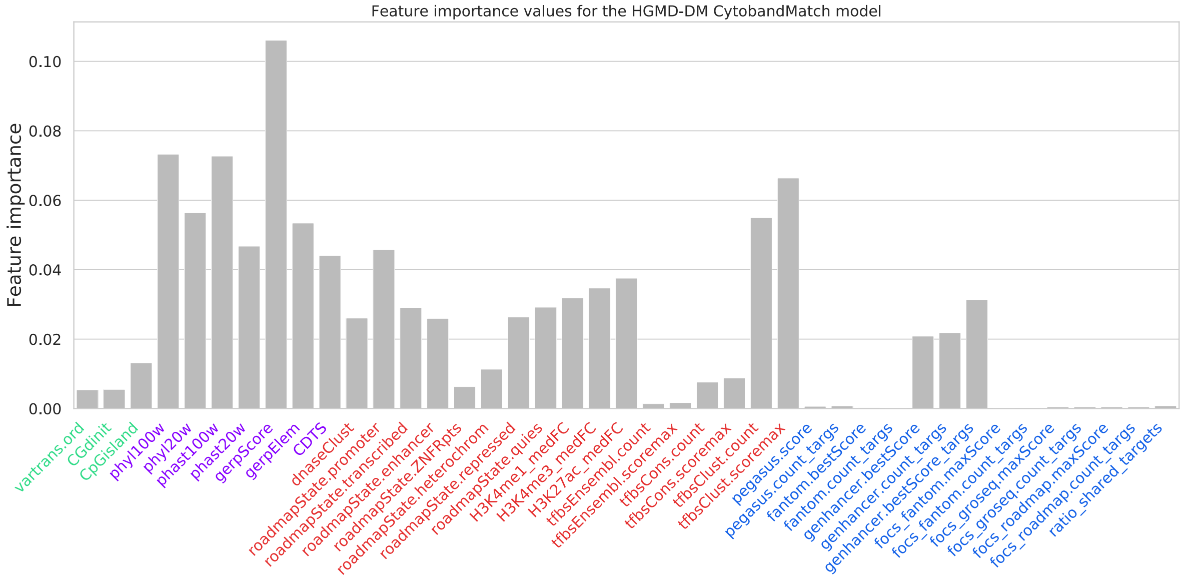


**Supplementary Figure 3.** Feature importance of the 41 features in the FINSURF Adjusted model. Features are listed on the x-axis and grouped in four categories: green for nucleotide sequence features, purple for evolutionary sequence conservation features, red for functional genomics features and blue for putative enhancer – promoter association features.

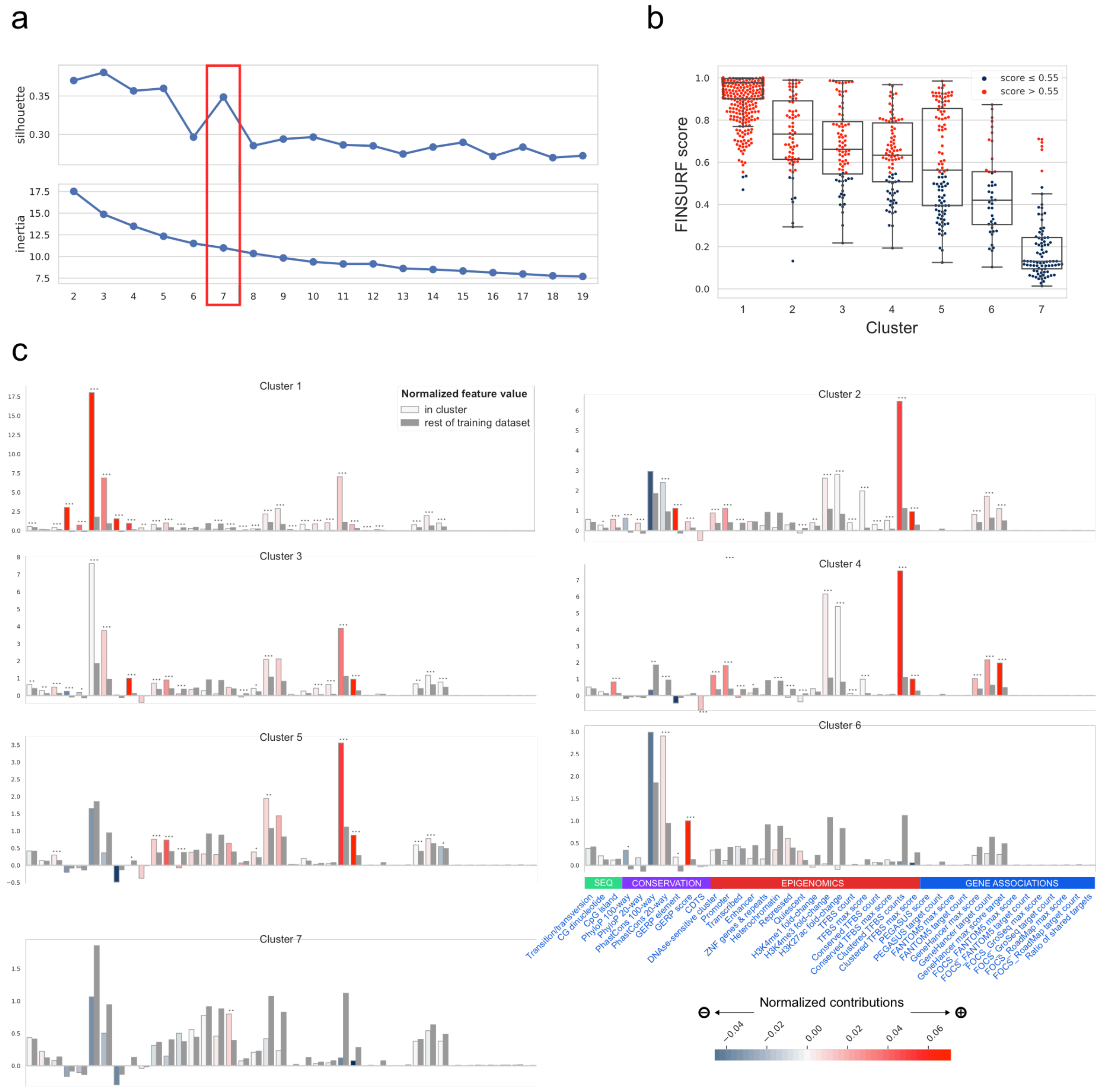


**Supplementary Figure 4**. Analysis of the 878 positive control variants of the FINSURF model through their feature contribution profiles. Contributions to the prediction score were calculated for each variant, and k-means clustering was applied. **a**. Different number of clusters were explored, and the finale number K=7 was selected from the joint minimization of the inertia and maximization of the silhouette. **b**. Distribution of scores of variants per cluster, colored by the predicted class when applying the optimal threshold of 0.51. **c**. Average functional profile of the 7 clusters. These profiles highlight how feature contributions relate to the raw values of the features, as processed by the random forests. For each cluster, the raw values were normalized, and averaged from the variants within the cluster (colored bars). Reference values (dark grey bars) correspond to the average from the rest of the training dataset (i.e. positive controls in other clusters and negative controls). The color of the bars is based on the average contributions of the features in each cluster.

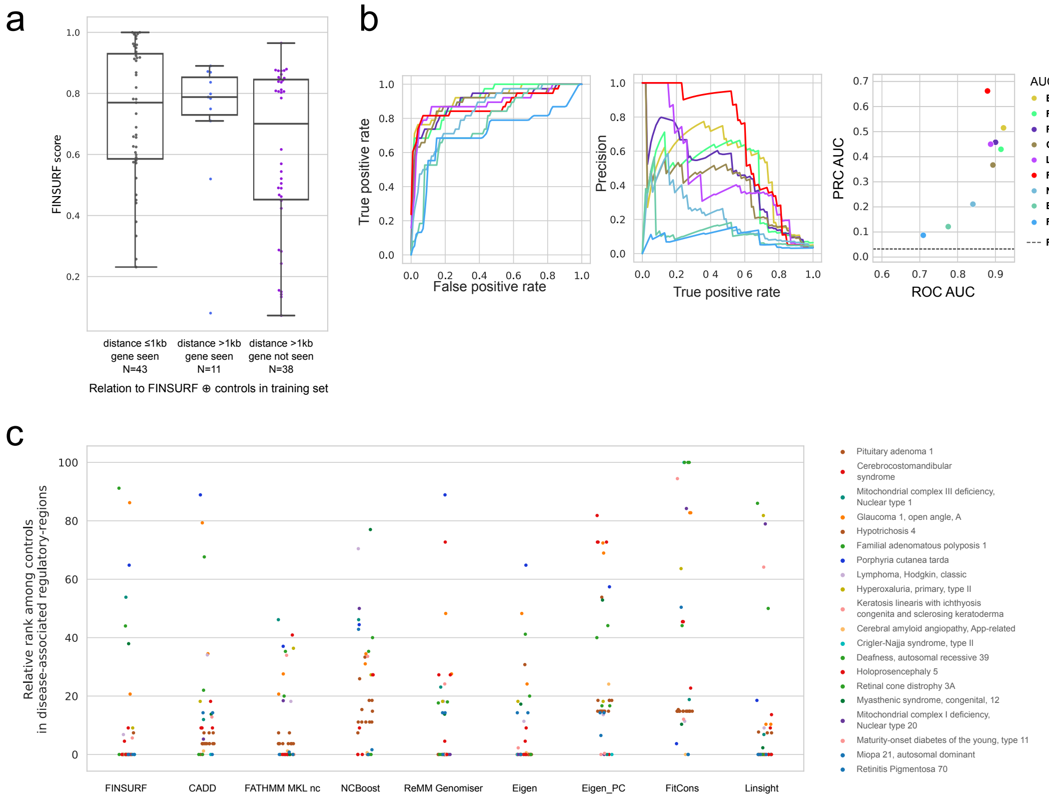


**Supplementary Figure 5.** Predicting highly pathogenic disease variant in genomic context. **a**. Box-plots representation and individual FINSURF score values for 92 known pathogenic variants from the Genomiser dataset but absent from the FINSURF training set. While 43 variants are very close (≤ 1kb) to one of the latter (left), the remaining 49 are sufficiently far (> 1kb) to be considered to overlap independent annotations. Of these a subset of 38 variants are also predicted to reside in a regulatory element targeting a gene that is not in the set of targets of variants in the FINSURF training set ("not seen"). Since the two subsets (N=11 and N=38) show largely overlapping score distributions, they were all used for classification performance in a genomic context. **b**. Classification performance comparisons between FINSURF and eight other methods on the set of 49 positive variants described in (a): left, ROC; middle, PRC; right, scatterplot of ROC versus PRC AUC values for graphs shown on the left and middle panels. Note that only 38 variants were eventually scored by all 9 methods and could be used for the analysis. The negative set used here were the 1,140 Platinum variants also residing in the same predicted regulatory variants as the 38 positives. **c**. Comparison of ranks for the 38 positive variants among the 1,140 negatives, based on scores attributed by FINSURF and eight other methods. Ranks were normalised between 0 and 100 for each method (y-axis). Colours represent different diseases. The graph shows that there is little correlation between variant ranks among the different methods.
